# The sequence dependent search mechanism of EcoRI

**DOI:** 10.1101/444042

**Authors:** S.C. Piatt, J.J. Loparo, A.C. Price

## Abstract

One-dimensional search is an essential step in DNA target recognition. Theoretical studies have suggested that the sequence dependence of one-dimensional diffusion can help resolve the competing demands of fast search and high target affinity, a conflict known as the speed-selectivity paradox. The resolution requires that the diffusion energy landscape is correlated with the underlying specific binding energies. In this work, we report observations of one-dimensional search by QD labeled EcoRI. Our data supports the view that proteins search DNA via rotation coupled sliding over a corrugated energy landscape. We observed that while EcoRI primarily slides along DNA at low salt concentrations, at higher concentrations its diffusion is a combination of sliding and hopping. We also observed long-lived pauses at genomic star sites which differ by a single nucleotide from the target sequence. To reconcile these observations with prior biochemical and structural data, we propose a model of search in which the protein slides over a sequence independent energy landscape during fast search, but rapidly interconverts with a “hemi-specific” binding mode in which a half site is probed. This half site interaction stabilizes the transition to a fully specific mode of binding which can then lead to target recognition.

## INTRODUCTION

Many site-specific DNA binding proteins perform one-dimensional diffusive scans after encountering non-specific DNA. This idea not only explains biochemical data from several systems (1–3), but has also been directly verified in many cases through single molecule tracking of LacI (4), Rad51 (5), hOGG1 (6), p53 (7), as well as EcoRV (8). Studies of p53 (9) and zinc finger proteins (10) show that mechanisms of one-dimensional search can vary, requiring distinct intermediates for rapid and accurate search. Type II restriction endonucleases (2REs) were among the first DNA binding proteins to show one-dimensional search (11) and remain ideal model systems for the study of DNA target search (12) as well as of site specific DNA cleavage (13). Although significant structural and biochemical data exist for this class of enzymes, the mechanisms of one-dimensional search by 2REs are poorly understood (14).

Two microscopic mechanisms have been proposed to contribute to one-dimensional search (15). In sliding, the protein remains in contact with the DNA helix, rotating as it moves. Protein translocation steps involve moving to an adjacent non-specific binding site without ion recondensation onto the DNA backbone, implying that the rate of diffusion should have little dependence on the salt concentration. Diffusion coefficients independent of salt concentration have been observed for hOGG1 (6) and T7 RNAP (16), consistent with a sliding mechanism. This mechanism can produce thorough searches, as every site in the protein’s path is visited. In hopping, the protein dissociates from the DNA, allowing ion recondensation. Due to recurrence (revisiting the DNA), there is a significant probability the protein will rebind at a nearby site. Since the hopping motion is free three-dimensional diffusion, this motion will also have a weak dependence on salt concentration. Hopping, in contrast to sliding, can produce highly *transparent* paths, i.e., although the displacement along the DNA may be large, only a small fraction of the non-specific sites will be probed (17). Due to the strong salt dependence of the non-specific off rate, the balance between sliding and hopping will depend strongly on salt, and one-dimensional searches, which combine sliding and hopping, will have salt dependent diffusion coefficients. Such salt dependent diffusion coefficients have been observed for EcoRV (8), implying both mechanisms contribute under the buffer conditions used in that study.

In order to maintain contact with a non-specific binding site, a protein translocating along DNA must rotate to follow the helical backbone. Theoretical analyses have shown that this significantly reduces the one-dimensional diffusion coefficient relative to the three-dimensional constant (18, 19). Single molecule tracking experiments have obtained one-dimensional diffusion coefficients several orders of magnitude (10^3^ – 10^4^) lower than the three-dimensional constants (20). However, the rotational coupling is insufficient to explain the magnitude of the observed reduction. The remaining reduction is attributed to energy barriers between non-specific sites and to randomness in the sliding energy surface, typically of the order of 1 – 2 k_B_T (6–8). In spite of the importance of the theory of rotational coupling to our understanding of one-dimensional search, direct observation of rotation during sliding has yet to be made.

Rapid search requires a relatively smooth energy landscape with low barriers to translocation. However, target recognition complexes typically show extensive structural rearrangements indicating substantial barriers to specific association (14). Clearly, testing each potential binding site using the recognition conformation would significantly slow one-dimensional search. This conflict between the need for high affinity specific recognition and rapid diffusion has been referred to as the speed-stability paradox (21). Although the existence of this paradox has been questioned (22), models have been developed to explain how proteins can overcome this conflict (21, 23). These models assume that the protein exists in at least two conformations: a search mode, expected to participate only weakly in sequence specific interactions and which can diffuse quickly along the DNA, and a recognition mode, in which the protein adopts a conformation close to its specific binding configuration and is therefore inconsistent with rapid sliding. A key component of these models is “kinetic preselection.” The sliding energy surface over which the search conformation diffuses is correlated with the highly sequence specific energy landscape of the recognition mode. Hence the diffusing protein spends longer periods of time at sites with higher probability of being the target, reducing the time wasted probing unproductive sites. Such models have been applied to both p53 (9) and zinc finger proteins (10).

Structures of 2REs in complex with DNA have provided insight into the nature of the intermediates involved in one-dimensional search. The recognition complex of EcoRI with cognate DNA demonstrates extensive specific interactions via a recognition loop which extends down into the major groove (24). While complexes of EcoRI with non-cognate DNA have remained elusive, the related endonucleases, BamHI and BstYI, have yielded to crystallographic analyses (25, 26). Both of these proteins approach DNA from the major groove and cleave dsDNA leaving 5′ overhangs, as EcoRI does. As in the case of EcoRI, the cognate recognition complexes show significant specific interactions between DNA and recognition loops that enter the major groove. In contrast, the structures of BamHI and BstYI in complex with non-cognate DNA (in both cases, the DNA binding site differed in a single base pair from the target sequence, known as a star site) show the proteins in a more open conformation, with quaternary rearrangements resulting in a wider DNA binding cleft. In the non-specific complex, BamHI binds DNA symmetrically and does not protrude into the major groove, making no base specific contacts. On the other hand, BstYI binds in an asymmetric manner, rotated to bring the recognition loop of one monomer into the major groove, where it can make base specific interactions with the cognate half site (a binding mode termed “hemi-specific” by the authors of ref. (26)). The opposing monomer, rotated away from the major groove, does not make any specific contacts with the non-cognate half site. Whether these two non-specific binding modes represent true intermediates along the one-dimensional search pathway remains an open question.

In this paper we report our measurements of one-dimensional search of EcoRI along non-specific DNA. Using TIRF imaging of a quantum dot labeled protein, we observed one dimensional diffusion as well as pausing of the endonuclease on flow stretched λ DNA. Using the sliding diffusion coefficient, we determined that the average energy barrier to translocation was similar to that measured for other DNA binding proteins. The diffusion coefficient increased with salt concentration demonstrating that the protein diffuses via sliding and hopping. Finally, we integrate our observations of pausing with existing biochemical and structural data on one-dimensional search by 2REs to propose a model in which the protein first identifies half cognate sites before the transition to a full recognition complex where the entire target sequence is probed.

## MATERIALS AND METHODS

### Data collection

The conjugation of antibody to quantum dots was carried out according the Thermofisher CLICK conjugation protocol. The purification of EcoRI^E111Q^ protein, the functionalization of glass coverslips and the preparation and labeling of biotinylated λ DNA with quantum dots were carried out as described in Graham et al. (27). Briefly, a flow cell comprised of a coverslip functionalized with a mixture of PEG and PEG-biotin and a linear channel 1.8 mm wide and 120 μm high was used. The surface of the flow cell was coated with streptavidin, washed, and incubated with EcoRI^E111Q^-QD-labeled λ DNA for ~10 min or until tethers could be clearly seen. During data collection, the flow cell was washed with 50 pM QD labeled protein in 10 mM Tris, pH 8.0, 30-150 mM NaCl, 0.5 mM Mg_2_Cl, 200 μg/mL BSA. Labeled proteins were imaged with a home-built through-objective total internal reflection fluorescence microscope using 532 nm excitation (Sapphire 488-50, Coherent, Santa Clara, CA). Details of the microscope and its alignment can be found in Graham et al. (27). Video data was collected at 30 fps for 120s for salt concentrations 70 mM and above, and for 1200s for salt concentrations 60 mM and below.

### Data analysis

The region of the DNAs free of specifically bound proteins were identified as regions of interest (ROIs) for nonspecific sliding events. Tracking of proteins in the ROIs was completed using the ImageJ plugin, Fiji TrackMate (https://imagej.net/TrackMate). Custom Python code was used to correct for finite extension of DNA, analyze mean-squared deviation (MSD) and perform linear fits to determine diffusion coefficients. Particles in ROIs with diffusion coefficients not statistically different from zero were categorized as paused. The mean distance of paused proteins from specifically bound EcoRIs was used to locate pause sites in the λ genome. To determine the drift speed of the proteins, the slope of the longitudinal trajectories was determined by linear least-squares regression. Drift speeds at each salt concentration from 30 mM to 90 mM were determined. Drift speed at 150 mM was not well determined due to the short interaction time.

The p-value for the observed distribution of pauses was calculated using a Monte Carlo algorithm that simulated randomly distributed sets of pause locations and compared these to the true genomic star site distribution. The genomic star site distribution was determined using the star sites GAATTT, GAAGTC, GAATTA, and GAACTC, which were the four most frequent star sites cleaved by EcoRI in a whole genome study (28). Randomly generated distributions that had a Pearson correlation coefficient of 0.844 (the observed correlation) or better with the genomic star site distribution were counted as successes. Out of 5×10^7^ simulations, only 16255 successes were counted, indicating a *p*-value of 3.3 × 10^−4^.

## RESULTS

To characterize the one dimensional search mechanisms of EcoRI, we imaged single QD-labeled, catalytically inactive EcoRI (EcoRI^E111Q^) interacting with flow-stretched λ DNAs (Fig 1). Lambda DNA contains five cognate EcoRI sites located in the second half of its genome. The first half of the genome contains approximately 21 kbp of DNA lacking cognate sites, and this half was tethered to the flow cell surface through the 5’ end. Nonspecific DNA interactions were analyzed by restricting analysis to events on this first half of the DNA.

**FIGURE 1.**
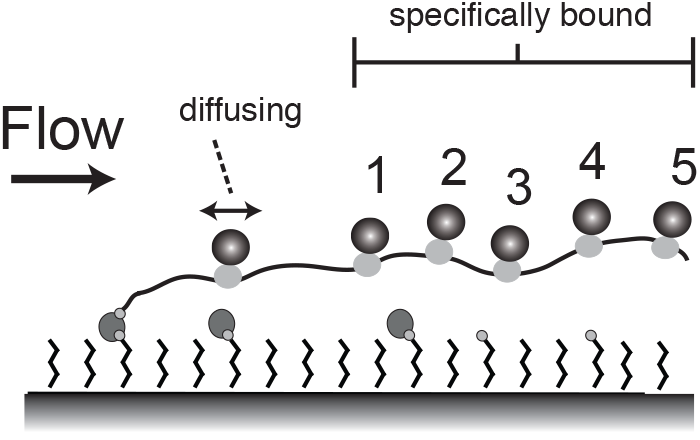
Experimental design. λ DNA is tethered to a glass surface via a biotin-streptavidin link and stretched by flow. QD labeled EcoRIE111Q is bound specifically to five approximately equally spaced cognate sites on free end of the molecule (to right in figure). Non-specifically bound EcoRI^E111Q^ (on left) is free to diffuse along the DNA.

We pre-incubated DNAs with QD-labeled EcoRIE111Q in order to label the five cognate sites prior to DNA tethering. The labeled cognate sites remained identifiable throughout data collection and served two important functions. First, they allowed for the rapid identification of tethered DNAs in a field of view. Second, they functioned as fiduciary markers that enabled us to determine the absolute location of any free EcoRI^E111Q^ interacting nonspecifically with the DNA.

We were able to observe non-specific binding of EcoRIE111Q to both doubly-tethered DNAs in the absence of flow as well as to singly-tethered DNAs elongated under flow. A relatively low concentration of labeled EcoRI (50 pM) was necessary to reduce background. Under zero flow conditions we had difficulty distinguishing nonspecifically bound proteins diffusing on the doubly tethered DNA from free proteins diffusing near the DNA. Applying low flow (25 μL/min) eliminated this background of freely diffusing protein. Therefore, the results we report here are from measurements of the singly-tethered DNAs under low flow conditions.

Accurate positions of the labeled proteins were determined by fitting Gaussian intensity profiles to the diffraction-limited images of EcoRI^E111Q^. Absolute locations in the λ genome were determined by measuring the distance to the labeled specific sites and correcting for the extension of the DNA. To determine the DNA extension, we first calculated the Weissenberg number (Wi). The Weissenberg number is a dimensionless parameter that completely characterizes the dynamics of singly tethered polymers under shear flow. Using our channel geometry and flow rate, we calculated Wi = 19. We previously showed that at Wi = 19 the extension of QD labeled λ DNA is 66% and is relatively constant from the tether point out to the second QD (29). We next experimentally measured the extension using position measurements of QDs 1 and 2 and found the extension to be 65%, in agreement with our first method. All the non-specific interactions we report in the current work were observed in the region between the tether point and the first QD, corresponding to a contour length of around 7 μm of DNA. All data are corrected for the uniform extension of 65%.

In many cases, we observed one-dimensional diffusion of EcoRI on λ DNA. An example of a diffusing trajectory is shown in Fig. 2A. To further analyze the motion, we calculated the mean squared deviation (MSD) of the longitudinal (along the DNA) coordinate as a function of the time interval. The MSDs (Fig. 3A) displayed a linear dependence on time with a small fast component with a rise time of a few milliseconds. The transverse coordinate showed constrained motion, consistent with a protein moving along a flow stretched DNA. The MSDs of the transverse coordinate (Fig. 3B) were independent of time except for a fast component with a rise time similar to the fast component in the longitudinal trajectories. The timescales (a few ms) and the amplitudes (~100 nm) of these fast components are due to the underlying fluctuations of DNAs extended in shear flow and are similar to values we have previously reported (29).

**FIGURE 2.**
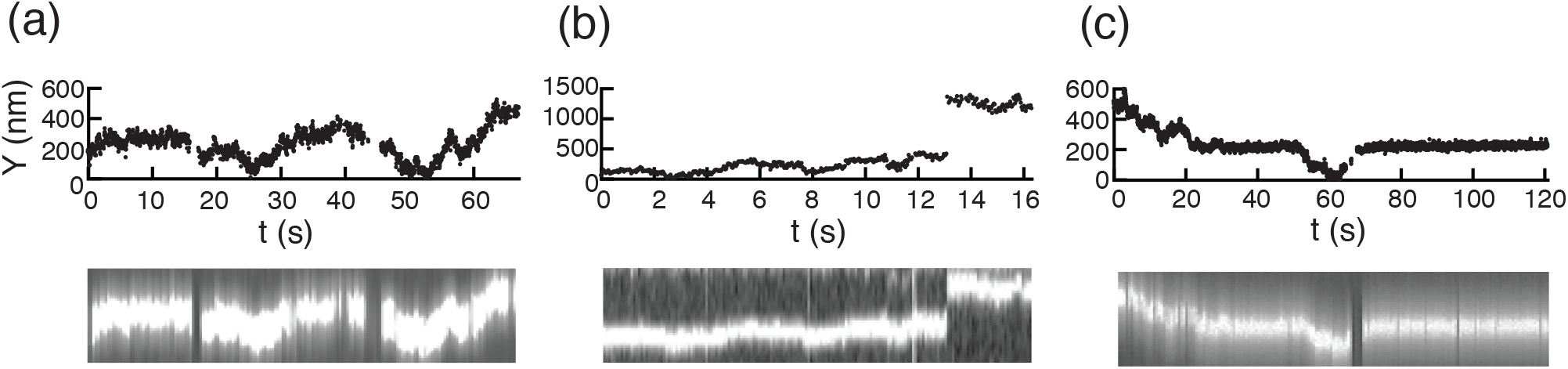
Kymographs and tracking of EcoRI^E111Q^ molecules diffusing on DNA. (a) A single continuous diffusion trajectory. (b) Two continuous trajectories connected by a jumping event. (c) A continuous diffusion trajectory interrupted by two pausing events.

**FIGURE 3.**
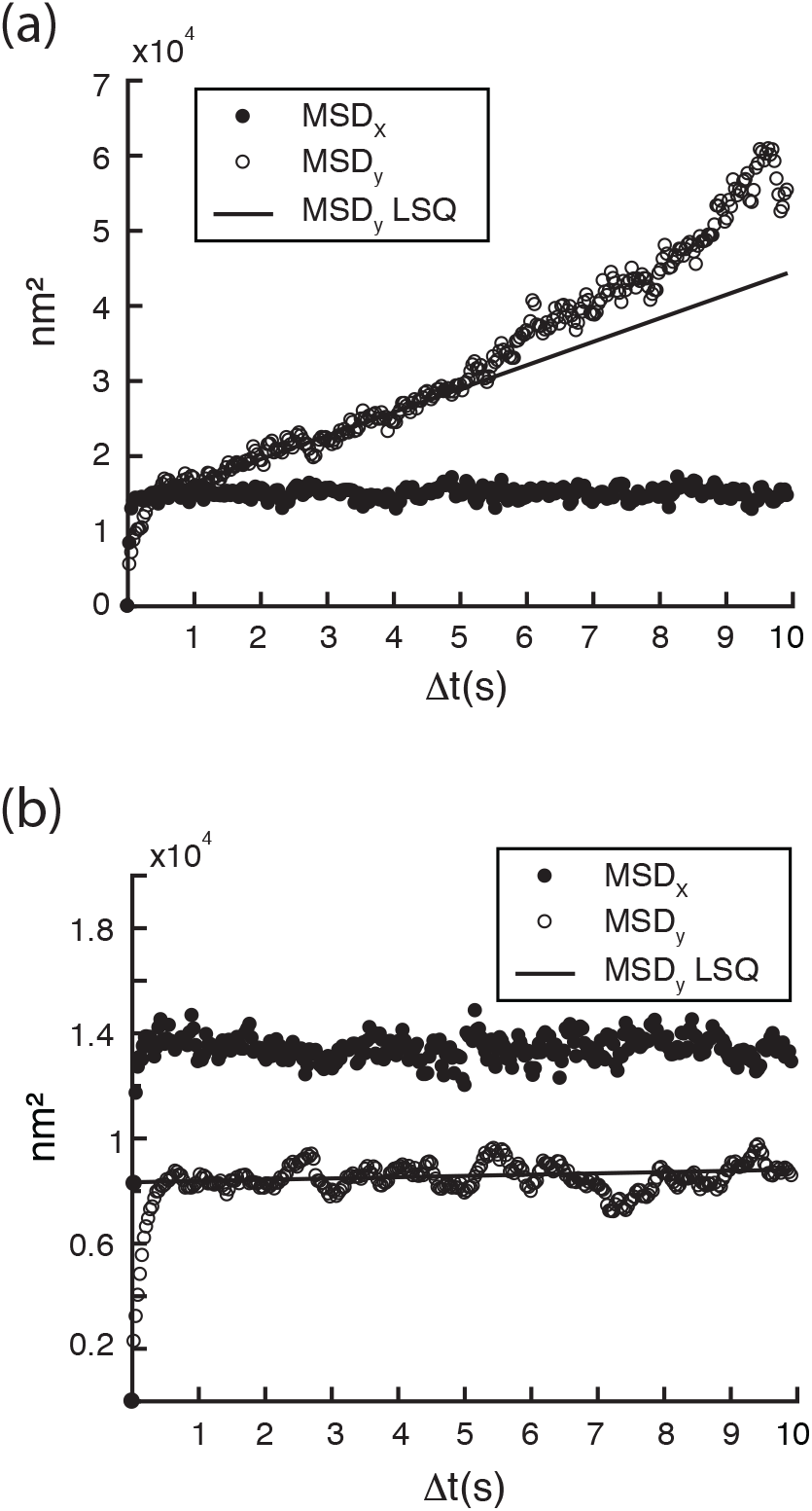
MSD plots of (a) freely-diffusing and (b) paused molecules in the transverse (x) and longitudinal (y) directions. In panel (a) the initial slope (t < 5 s) of the MSD in the longitudinal coordinate was used to determine diffusion coefficient along the DNA. The initial fast rise in the MSDs is due to the dynamic fluctuations in the DNA.

To determine how much effect the flow in our experiment had on our results, we measured the average drift speed of the protein along the DNA. We found a salt dependent drift speed that varied from ~0.5 nm/s at 30 mM NaCl to ~2 nm/s above 70 mM NaCl. This drift accounts for 5% to 15% of average length of DNA scanned per one-dimensional search, indicating that the drift did not play a major role in the motion in our experiments.

We determined one-dimensional diffusion coefficients from individual trajectories of EcoRI position using linear fits to the MSD curves of the longitudinal coordinates (Fig. 3A). To further minimize the effect of the bias due to flow, we fit the MSDs to only the first few seconds (3 – 5s) to determine the initial slope. In order to determine the relative contributions to the diffusion from sliding and hopping, we measured how the one-dimensional diffusion coefficient varied with salt concentration (Fig. 5). The diffusion coefficient was independent of salt concentration from 30 mM to 70 mM, with a mean of 1.3×10^−3^ μm^2^/s, implying that sliding was the predominant mechanism of search. Consistent with a sliding based search, we observed very few jumps in longitudinal position (5 out of 208 events, see example in Fig. 2B). However, above 70 mM NaCl, the diffusion coefficient increased, reaching more than three times the low salt value at 150 mM, implying that hopping contributes more at these higher salt concentrations.

**FIGURE 4.**
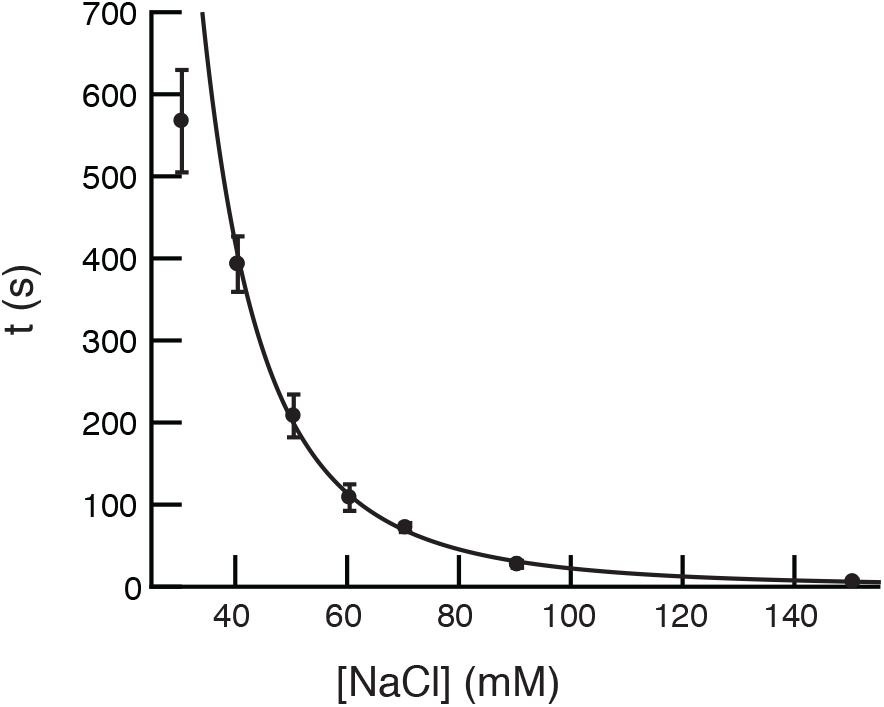
Mean dwell times of diffusing EcoRI^E111Q^ versus salt concentration. The dwell time was determined as the time from initial appearance on the DNA until dissociation, indicated by disappearance of the fluorescently labeled protein (114 events). The solid curve is a fit to a power law (exponent = 3.2 ± 0.1) Error bars represent the standard error of the mean.

**FIGURE 5.**
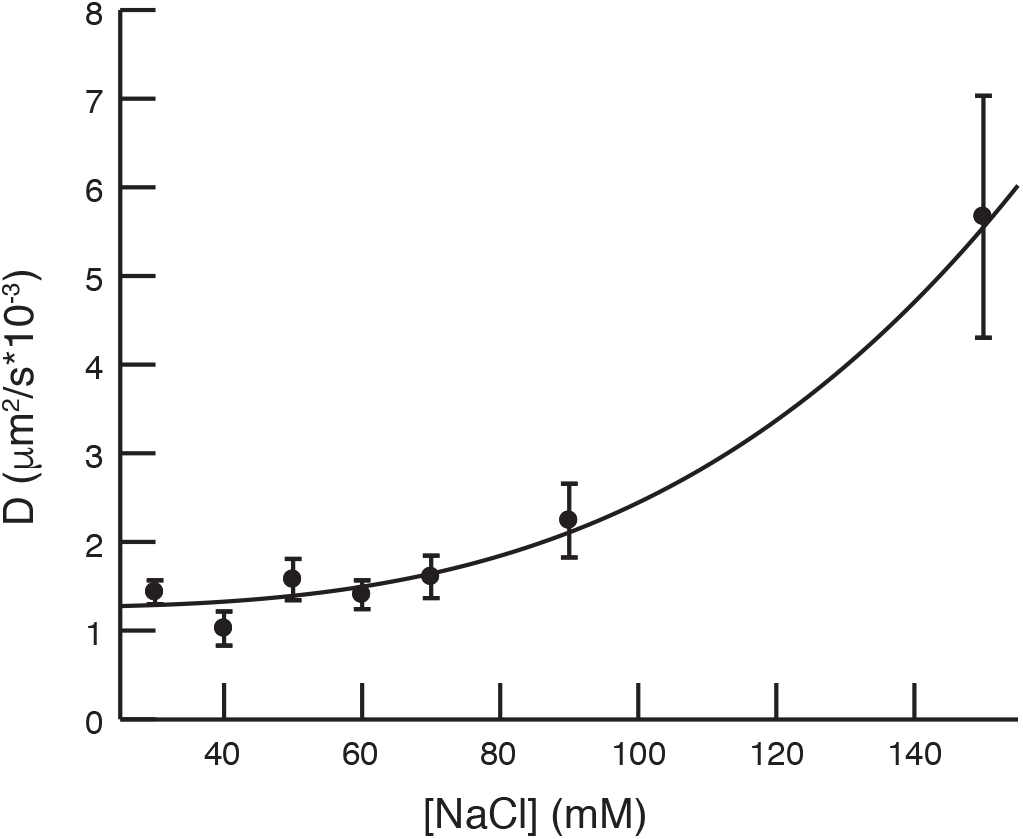
One-dimensional diffusion coefficient versus salt concentration. Shown is the mean of the diffusion coefficients at each salt concentration as determined by linear fits to MSDs (114 events). The solid curve is a fit to Eq. 1. Error bars represent the standard error of the mean.

The mean dwell times of the one-dimensional diffusive interactions show a strong dependence on salt concentration (Fig. 4). Assuming the dissociation reaction follows the law of mass action, we fit the measured dwell times to a power law of the form A[Na^+^]^−q^, where A and q are fitting parameters. The exponent fits to a value of *q* = 3.2 ± 0.1, and can be interpreted as the number of counter ions released from the DNA upon non-specific binding of the protein.

We also determined the one-dimensional scan range (defined as the maximum minus minimum observed longitudinal coordinate for a single diffusion event) for each salt concentration. Figure 6 shows this range as well as the expected RMS displacement for each trajectory as calculated from the measured diffusion coefficients and dwell times. At [NaCl] < 70 mM, the range decreases strongly as salt increases due to the dependence of the dwell time on salt. However, at higher salt the scan range and RMS displacement only depend weakly on salt, as the reduced dwell time is compensated for by the increase in the diffusion coefficient.

**FIGURE 6.**
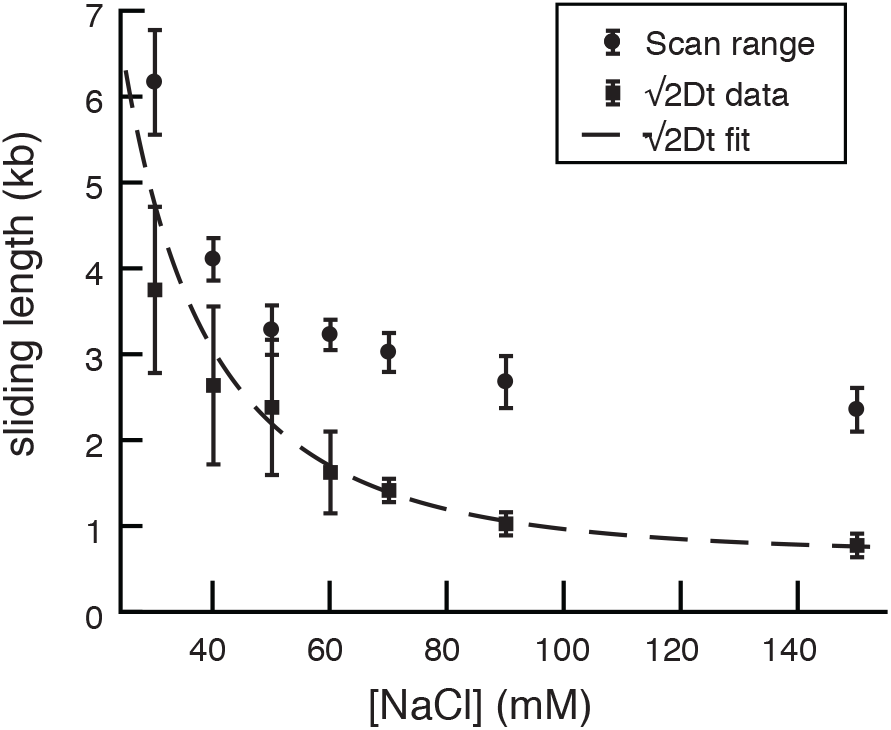
Mean scan range of diffusion events versus salt concentration. The scan range was calculated as the maximum length scanned in the one dimensional diffusion event (i.e., the maximum minus the minimum longitudinal coordinate in a single diffusion trajectory). Also plotted is the diffusion length, defined as the predicted RMS displacement due solely to diffusion 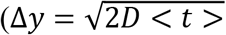,where <t> is the mean dwell time from Fig. 4 and D is the measured diffusion coefficient from Fig. 5). The dashed curve is the predicted diffusion length from Eq. 1.

A significant number of trajectories (45%) showed pausing of EcoRI during one-dimensional diffusion. Some trajectories showed the enzyme diffusing one dimensionally to the pause site where it stalled upon reaching the site. In other trajectories, the protein could be seen leaving the pause site, diffusing away and then returning to pause once again at the same (or indistinguishable nearby) site (Fig. 2C). In yet other cases, the non-specifically bound protein did not display any diffusion along the DNA during the entire data collection. Using the labeled cognate sites for reference, we were able to map the genomic locations of these pause sites and determine that many of them clustered into distinct regions of the DNA, indicating that EcoRI pauses at several specific non-cognate sites in the λ genome. These observations led us to consider whether these pause sites could be EcoRI star sites. Star sites vary by a single base pair from the cognate sequence. High throughput sequencing methods have characterized star activity in whole bacterial genomes (28). We mapped out the location of the four most prevalent EcoRI star sites identified in ref. (28) and compared these to our observed pause sites (see Fig. 7). The high correlation between the two distributions (*p* = 3.3 × 10^−4^) indicates that EcoRI pauses preferentially at star sites.

**FIGURE 7.**
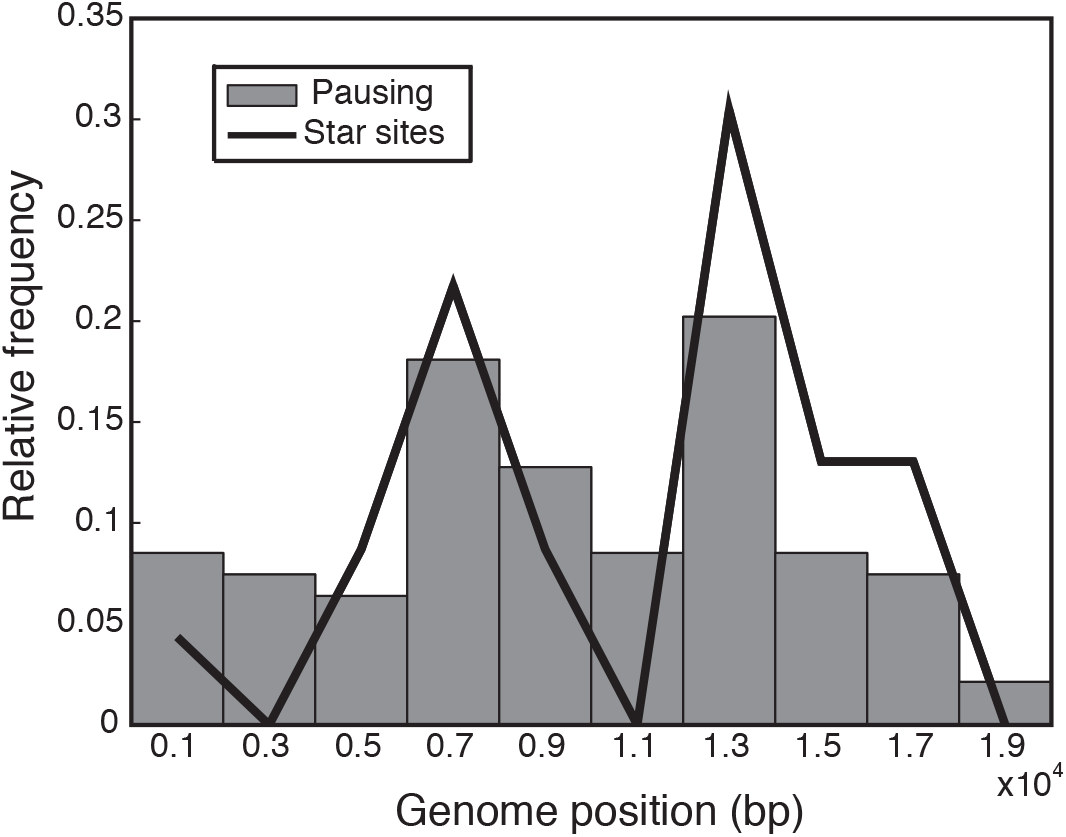
Comparison of observed pause sites and star sites. Relative frequency of pausing (94 events, shown in columns) and genomic star sites (solid lines) show high correlation (0.83). The probability of obtaining the observed correlation assuming randomly distributed pause sites (p-value) is 3.5×10^−4^.

## DISCUSSION

### The interaction of EcoRI with non-specific DNA

In the presence of a single species of counter ion, the non-specific off-rate is proportional to a power of the ion concentration with the exponent equal to the number of cations displaced upon protein binding. The crystal structure of EcoRI in complex with cognate DNA shows that the protein completely covers the six recognition bases, implying ~12 phosphate groups are blocked by the EcoRI footprint (24). If we assume the footprint of the non-specific complex to be similar, we expect a release of ~9 ions upon dissociation of the non-specific complex. (This assumes an occupation of 0.76 of the phosphate groups by counter ions (30).)

Our data shows that the non-specific off rate scales with salt concentration with an exponent of 3.2 ±0.1 (Fig. 4), significantly less than the expected value based on the footprint of the cognate complex (~9). One complication of our analysis is that the interactions we observed are most likely made up of multiple cycles of sliding and hopping. This is because small hops cannot be distinguished from the dynamic fluctuations in the DNA. The total dwell time we measure is *t* = *N*(*τ_S_* + *τ_H_*), where *N* is the number of cycles, *τ_S_* is the mean time per sliding event, and *τ_H_* is the mean time per hop. The number of hopping cycles will not depend strongly on salt, since it only depends on the probability of recurrence once the protein has dissociated, a probability which is determined by the statistics of the three-dimensional diffusion. In addition, the mean time per hop is expected to be very small compared to the time in sliding, as it is controlled by the three-dimensional diffusion coefficient (15 μm^2^/s from ref (29)), which is significantly greater than that of the one-dimensional diffusion. Therefore, the scaling exponent we determined should reflect that of the intrinsic off rate from the sliding (non-specifically bound) state.

Recent biochemical experiments have measured the salt dependence of the dissociation of EcoRI and derived an exponent of 5.8 ±0.2, higher than our observed result, but still smaller than the expected footprint (31). Our direct observation of the interaction allows us to exclude interactions with pausing, which was not addressed in the biochemical study. Our results are consistent with a loose non-cognate complex that does not disrupt as many counter ion-phosphate interactions as the cognate complex.

### Sliding and hopping both contribute to 1D search

Due to the strong salt dependence of non-specific dissociation, the relative contributions of sliding and hoping to one-dimensional diffusion will also depend on salt. Due to the faster hopping motion, the diffusion coefficient should increase as the sliding contribution decreases. In the case of EcoRI, our data shows two limits of behavior. Below 70 mM NaCl, sliding dominated diffusion results in a diffusion coefficient that is largely independent of salt. However, at higher salt concentrations, the relative contribution of hopping to sliding increases thus resulting in the diffusion coefficient increasing with salt concentration. This behavior of EcoRI is in contrast to other proteins that have not shown this transition from predominantly sliding to mixed sliding/hopping diffusion. For example, the diffusion coefficient of EcoRV (8) was observed to increase with salt over the entire range examined (10 – 60 mM NaCl). In the cases of T7 RNAP (salt range 0 – 50 mM NaCl, ref. (16)) and hOGG1 (salt range 10mM – 100mM NaCl, ref. (6)), no salt dependence was observed.

When both mechanisms are present, the one dimensional diffusion coefficient *D_1_* will be a weighted average of two diffusion coefficients (32):

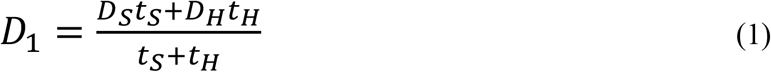

In this equation, *D_S_* and *D_H_* are the sliding and hopping diffusion coefficients, and *t_S_* and *t_H_* are the mean total time spent in sliding and hopping. The three factors *t_H_*, *D_S_* and *D_H_* (≅ *D_3_*, the three dimensional diffusion coefficient) will only depend weakly on salt concentration. The only strong dependency on salt then is through the *t_S_* term. When *D_S_t_S_* >> *D_H_t_H_* (at low salt), *D_1_* is equal to *D_S_*. At high salt, when *t_S_* << *t_H_*, *D_1_* will again be independent of salt and will be equal to *D_H_* ≅ *D_3_*. This second limit may be unobservable, as it may only apply at extremely high salt. At intermediate concentrations, *D_1_* will depend strongly on salt. Assuming the residence time of the protein on non-specific DNA follows a power law of the form t ~ [Na^+^]^−q^, the diffusion coefficient will scale with salt concentration ~[salt]^q^ in this intermediate regime.

To determine quantitatively the separate contributions of sliding and hopping, we have fit the above equation to our data (see Fig. 5). Using the fit from the residence times for the *t_S_* term, and assuming *D_H_* = *D_3_*, there are only two free parameters, *D_S_* and *t_H_*. The constant value for *D_1_* that we observe at low salt determines *D_S_* = (1.26±0.09) × 10^−3^ μm^2^/s and the critical salt value when *D_1_* begins to increase determines *t_H_* = 1.4 ±0.4 ms. This later value implies that EcoRI spends ~1.4 ms in hopping during each one dimensional scan (excluding pauses). Our data does not allow us to determine the number of hops per scan. However, the fit to the model determines that sliding and hopping make equal contributions to the mean squared displacement at a salt concentration of ~110 mM, where the total RMS displacement due to diffusion is ~1000 bp for each one-dimensional scan.

As the salt concentration is increased above the critical value, the hopping distribution will remain relatively constant as the sliding length (the length of DNA probed during single sliding events) reduces. This will lead to an increase in the transparency, as the hops will remain the same size, but the regions of DNA probed during the sliding phase will decrease, eventually becoming much smaller than the hop step size. The length of DNA scanned per encounter will remain relatively constant in this limit. This is illustrated in our data (Fig. 6) that shows that the scan range depends weakly on salt above 70 mM, where the hopping is predicted to play a more significant role in diffusion. This increase in transparency has a major effect on the effectiveness of the one-dimensional scan. In contrast to EcoRV, whose critical salt concentration seems to be less than 10 mM (8), the critical concentration for EcoRI is closer to physiological salt concentrations due to the longer residence time of the non-specifically bound EcoRI. This explains prior biochemical data that suggests at 50 mM NaCl, EcoRI effectively searches all intervening DNA when translocating between two sites (33). At this salt concentration, transparency is low.

A significant amount of biochemical characterization of the one-dimensional search by DNA binding proteins has been carried out at low salt. At high salt, higher transparency will lead to reductions in efficiency. At low salt, DNA binding proteins can use a “single hit” approach, where a single non-specific encounter can lead to target acquisition. This can lead to specific on rates proportional to DNA length, as has been observed for 2REs (2, 11). It is unlikely that this limit is valid under physiological conditions.

### Rotational coupling and the energy landscape of sliding

Proteins that slide on DNA rotate following the helical backbone as they translocate to maintain contact with the non-specific binding site. This rotational coupling is a major contributor to the sliding diffusion coefficient, significantly reducing *D_S_* compared to the free three-dimensional diffusion coefficient, *D_3_*. In addition, thermodynamic energy barriers to single base pair translocation can reduce *D_S_* further. It has also been shown that disorder (or roughness) in the energy barriers, as well as randomness in the depths of the local non-specific binding energy wells, can further slow diffusion (21, 34).

Our data allows us to unequivocally determine the sliding diffusion coefficient *D_S_*, allowing us to apply theories of rotation coupled sliding to our measured value of (1.26±0.09)×10^−3^ μm^2^/s. The theory of rotation coupled sliding states that the one dimensional diffusion coefficient for a sliding mechanism in the absence of energy barriers is (19)

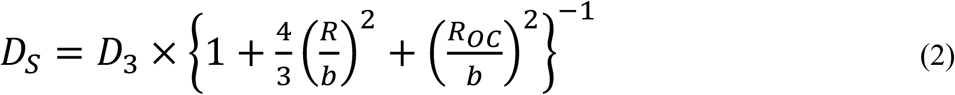

In this expression, *R* is the hydrodynamic radius of the protein, *R_OC_* is the distance of the center of the protein to the central axis of the DNA, and *b* is the pitch of the DNA (0.54 nm/rad). We can use our previous measurement of the hydrodynamic radius of the QD labeled EcoRI (13.7±0.4 nm, ref. (29)), and estimate *R_OC_* ≅ 13.7 ±1.0 nm. This leads to a reduction factor of 1500×(±9%) due solely to the rotation coupling as calculated using Eq. 2. The observed reduction factor is 15/0.00126 = 11900×(±10%), which is significantly greater than what rotational coupling would predict. The further reduction (11900/1500 = 7.9×) can be explained by the presence of thermodynamic energy barriers separating adjacent non-specific binding sites. Assuming the reduction follows an Arrhenius form (~exp(-ε/k_B_T), where ε is the activation energy), we determine a characteristic energy barrier of 2.1 ±0.4 k_B_T, a number comparable to that found for other DNA binding proteins (6–8). Roughness of the energy landscape (randomness in the activation energies), could be another factor in reduction of the diffusion coefficient. In a rough landscape, the average energy barrier will be less than the 2.1 k_B_T we determine here. The additional reduction in the diffusion coefficient will arise from randomness in the binding energies and barrier heights. However, our data does not allow us to independently determine the roughness parameter.

### Sequence dependent pausing during one dimensional search

Sequence dependent pausing of DNA binding proteins during one dimensional target search is poorly characterized. Kinetic data on association rates of EcoRI using DNAs containing star sites has been interpreted as implying that EcoRI pauses at these star sites for up to 20s in 50 mM NaCl (33). Through direct imaging of EcoRI we observe that it can remain bound to non-specific sites for many minutes. These long pauses likely result from multiple unobserved small excursions and returns to the star site. We can assume the minimum distance the protein must diffuse before we can identify an excursion is one half the RMSD dynamic fluctuations in the DNA. This is ~200 bp as determined from the y-intercept in Fig. 3B. This value is consistent with the track shown in Fig. 2C, which shows a just detectable excursion from the pause site at ~30 s with an amplitude of ~200 bp. Assuming the protein making the excursion is only sliding and starts from a non-specific binding site displaced a singe basepair from the pause site, this implies an escape probability of ~0.5%. The observed pauses are then composed of multiple “micropause” events, each linked by a short excursion and return. In the biochemical experiments in ref. (33), the cognate site was less than ten base pairs from the star site, and hence recurrence before specific association with the target site was much less likely. This picture agrees with the very long duration of the pauses we observed, many in excess of 20 minutes.

A model of the paused state has been proposed based on crystal structures of BstYI (26). In complex with star DNA, only one monomer of BstYI reaches into the major groove to make specific contacts, while the other is rotated out where it only makes non-specific interactions. This mode of binding (termed “hemi-specific” by the authors of ref. (26)) suggests widely distributed pauses which should occur at all sites that contain a single half cognate site. The observed distribution of EcoRI pausing shows enhancement at star sites. To reach the cognate-like interaction points in the non-cognate half of the star site, EcoRI must adopt a more closed conformation, similar to the specific binding mode.

### A model of 1D search by EcoRI

The above considerations suggest a model for sliding in which the protein adopts one of two types of interactions with non-cognate DNA (see Fig. 8). In the non-specific mode, the protein is positioned in a symmetric manner in the major groove and makes no specific contacts, similar to what has been observed for BamHI (25). Such an interaction is consistent with rapid translocation to neighboring sites and would lead to a uniform binding energy surface. In the hemi-specific mode (analogous to the BstYI-star structure), the protein is rotated, bringing one monomer into contact with the major groove where it has access to specific contacts in a potential half site. Importantly, the non-specific and hemi-specific modes can rapidly interconvert, as they are principally related by a rigid body rotation of the protein. Short pauses occur when target like interactions with the probed half site stabilize the protein-DNA complex. From this short pause state, further conformational changes (which can involve both closing of the protein dimer and bending and opening of the major groove) occur which bring the opposite binding site in the protein into contact with the remaining half site. Cognate interactions in the second half site will necessarily further stabilize the recognition complex, leading to a longer duration pause. It is these longer, sequence dependent pause states which we observe, as the shorter pauses are likely too fast to detect with our time resolution (~30ms). Once stabilized, the specific mode of binding can then lead to target recognition and subsequent hydrolysis of the phosphodiester bond.

**FIGURE 8.**
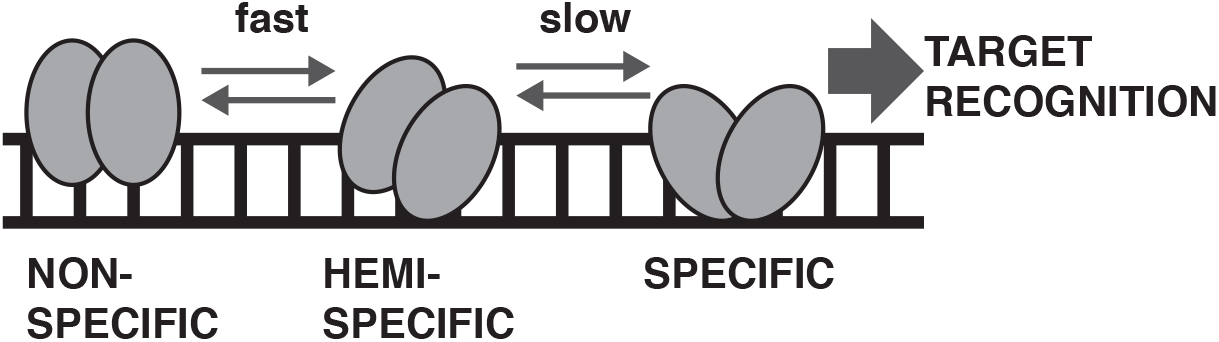
A model for one-dimensional search of DNA. In one-dimensional search, the protein rapidly intraconverts between the non-specific binding mode and the hemi-specific mode as it slides. Slower transitions to the specific mode of binding (which requires substantial conformational change) are more likely when cognate like interactions in a half site stabilize the hemi-specific binding mode. Target recognition, and subsequent catalysis, is only possible after the specific mode of binding is achieved.

In contrast to models which require kinetic preselection, this picture implies a relatively uniform non-specific energy landscape. In this way, the protein can rapidly search non-specific DNA and limit the occurrence of the specific binding mode to sites that are at least “half right.” This suggests a resolution to the speed/stability paradox similar to the resolution of Levinthal’s paradox of protein folding, in which the protein does not sample all possible folded conformations, but proceeds through a specific set of intermediate states. The short pause state (due to hemi-specific binding) as well as the longer pause state (with its more significant conformational changes) are then necessary intermediates which must occur before the formation of the protein-target recognition complex.

## AUTHOR CONTRIBUTIONS

SCP performed experiments, analyzed data and wrote paper. JJL designed research, wrote paper. ACP designed research, analyzed data, wrote paper.

## ACKNOWLEDGEMENTS

We would like to thank HyeongJun Kim for assistance in preparing the quantum dot labeled protein. Support for this work also comes from a National Institutes of Health grant R01GM115487 (to JJL) and the National Science Foundation RUI Award MCB-1715317 (to ACP).

## REFERENCES

1. Winter, R. B., O. G. Berg, and P. H. Von Hippel. 1981. Diffusion-driven mechanisms of protein translocation on nucleic acids. 3. The Escherichia coli lac repressor-operator interaction: kinetic measurements and conclusions. Biochemistry 20:6961–6977.

2. Terry, B., W. Jack, and P. Modrich. 1985. Facilitated diffusion during catalysis by EcoRI endonuclease. Nonspecific interactions in EcoRI catalysis. Journal of Biological Chemistry 260:13130–13137.

3. Gowers, D. M., G. G. Wilson, and S. E. Halford. 2005. Measurement of the contributions of 1D and 3D pathways to the translocation of a protein along DNA. Proceedings of the National Academy of Sciences of the United States of America 102:15883–15888.

4. Wang, Y., R. H. Austin, and E. C. Cox. 2006. Single molecule measurements of repressor protein 1D diffusion on DNA. Physical review letters 97:048302.

5. Granéli, A., C. C. Yeykal, R. B. Robertson, and E. C. Greene. 2006. Long-distance lateral diffusion of human Rad51 on double-stranded DNA. Proceedings of the National Academy of Sciences of the United States of America 103:1221–1226.

6. Blainey, P. C., A. M. van Oijen, A. Banerjee, G. L. Verdine, and X. S. Xie. 2006. A base-excision DNA-repair protein finds intrahelical lesion bases by fast sliding in contact with DNA. Proceedings of the National Academy of Sciences 103:5752–5757.

7. Tafvizi, A., F. Huang, J. S. Leith, A. R. Fersht, L. A. Mirny, and A. M. Van Oijen. 2008. Tumor suppressor p53 slides on DNA with low friction and high stability. Biophysical journal 95:L01–L03.

8. Bonnet, I., A. Biebricher, P.-L. Porte, C. Loverdo, O. Bénichou, R. Voituriez, C. Escude, W. Wende, A. Pingoud, and P. Desbiolles. 2008. Sliding and jumping of single EcoRV restriction enzymes on non-cognate DNA. Nucleic acids research 36:4118–4127.

9. Tafvizi, A., F. Huang, A. R. Fersht, L. A. Mirny, and A. M. van Oijen. 2011. A single-molecule characterization of p53 search on DNA. Proceedings of the National Academy of Sciences 108:563–568.

10. Iwahara, J., and Y. Levy. 2013. Speed-stability paradox in DNA-scanning by zinc-finger proteins. Transcription 4:58–61.

11. Ehbrecht, H.-J., A. Pingoud, C. Urbanke, G. Maass, and C. Gualerzi. 1985. Linear diffusion of restriction endonucleases on DNA. Journal of Biological Chemistry 260:6160–6166.

12. van den Broek, B., M. A. Lomholt, S.-M. Kalisch, R. Metzler, and G. J. Wuite. 2008. How DNA coiling enhances target localization by proteins. Proceedings of the National Academy of Sciences 105:15738–15742.

13. Gambino, S., B. Mousley, L. Cathcart, J. Winship, J. J. Loparo, and A. C. Price. 2016. A single molecule assay for measuring site-specific DNA cleavage. Analytical Biochemistry 495:3–5.

14. Pingoud, A., M. Fuxreiter, V. Pingoud, and W. Wende. 2005. Type II restriction endonucleases: structure and mechanism. Cellular and molecular life sciences 62:685–707.

15. Berg, O. G., R. B. Winter, and P. H. Von Hippel. 1981. Diffusion-driven mechanisms of protein translocation on nucleic acids. 1. Models and theory. Biochemistry 20:6929–6948.

16. Kim, J. H., and R. G. Larson. 2007. Single-molecule analysis of 1D diffusion and transcription elongation of T7 RNA polymerase along individual stretched DNA molecules. Nucleic acids research 35:3848–3858.

17. Redner, S. 2001. A guide to first-passage processes. Cambridge University Press.

18. Schurr, J. M. 1979. The one-dimensional diffusion coefficient of proteins absorbed on DNA. Hydrodynamic considerations. Biophysical chemistry 9:413–414.

19. Bagchi, B., P. C. Blainey, and X. S. Xie. 2008. Diffusion constant of a nonspecifically bound protein undergoing curvilinear motion along DNA. The Journal of Physical Chemistry B 112:6282–6284.

20. Blainey, P. C., G. Luo, S. Kou, W. F. Mangel, G. L. Verdine, B. Bagchi, and X. S. Xie. 2009. Nonspecifically bound proteins spin while diffusing along DNA. Nature Structural and Molecular Biology 16:1224.

21. Slutsky, M., and L. A. Mirny. 2004. Kinetics of protein-DNA interaction: facilitated target location in sequence-dependent potential. Biophysical journal 87:4021–4035.

22. Veksler, A., and A. B. Kolomeisky. 2013. Speed-selectivity paradox in the protein search for targets on DNA: is it real or not? The Journal of Physical Chemistry B 117:12695–12701.

23. Yu, S., S. Wang, and R. G. Larson. 2013. Proteins searching for their target on DNA by one-dimensional diffusion: overcoming the “speed-stability” paradox. Journal of biological physics 39:565–586.

24. Kim, Y., J. C. Grable, R. Love, P. J. Greene, and J. M. Rosenberg. 1990. Refinement of Eco RI endonuclease crystal structure: a revised protein chain tracing. Science 249:1307–1309.

25. Viadiu, H., and A. K. Aggarwal. 2000. Structure of BamHI bound to nonspecific DNA: a model for DNA sliding. Molecular cell 5:889–895.

26. Townson, S. A., J. C. Samuelson, Y. Bao, S.-y. Xu, and A. K. Aggarwal. 2007. BstYI bound to noncognate DNA reveals a “hemispecific” complex: implications for DNA scanning. Structure 15:449–459.

27. Graham, T. G., X. Wang, D. Song, C. M. Etson, A. M. van Oijen, D. Z. Rudner, and J. J. Loparo. 2014. ParB spreading requires DNA bridging. Genes & development 28:1228–1238.

28. Kamps-Hughes, N., A. Quimby, Z. Zhu, and E. A. Johnson. 2013. Massively parallel characterization of restriction endonucleases. Nucleic acids research 41:e119–e119.

29. Price, Allen C., Kevin R. Pilkiewicz, Thomas G. W. Graham, D. Song, Joel D. Eaves, and Joseph J. Loparo. 2015. DNA Motion Capture Reveals the Mechanical Properties of DNA at the Mesoscale. Biophysical journal 108:2532–2540.

30. Lohman, T. M., and P. H. von Hippel. 1986. Kinetics of Protein-Nucleic Acid Interactions: Use of Salt Effects to Probe Mechanisms of Interactio. CRC critical reviews in biochemistry 19:191–245.

31. Sidorova, N. Y., T. Scott, and D. C. Rau. 2013. DNA concentration-dependent dissociation of EcoRI: direct transfer or reaction during hopping. Biophysical journal 104:1296–1303.

32. DeSantis, M. C., J.-L. Li, and Y. Wang. 2011. Protein sliding and hopping kinetics on DNA. Physical Review E 83:021907.

33. Jeltsch, A., J. Alves, H. Wolfes, G. Maass, and A. Pingoud. 1994. Pausing of the restriction endonuclease EcoRI during linear diffusion on DNA. Biochemistry 33:10215–10219.

34. Hu, T., and B. Shklovskii. 2006. How does a protein search for the specific site on DNA: the role of disorder. Physical Review E 74:021903.

